# Spotsweeper-py: spatially-aware quality control metrics for spatial omics data in the Python ecosystem

**DOI:** 10.64898/2025.12.06.692760

**Authors:** Xingyi Chen, Michael Totty, Stephanie C. Hicks

**Affiliations:** Department of Applied Math and Statistics, Johns Hopkins University, Baltimore, MD, USA; Department of Biostatistics, Johns Hopkins Bloomberg School of Public Health, Baltimore, MD, USA; Department of Biomedical Engineering, Johns Hopkins School of Medicine, Baltimore, MD, USA; Center for Imaging Science, Johns Hopkins University, Baltimore MD 21218, USA; Johns Hopkins Kavli Neuroscience Discovery Institute, Baltimore, MD, USA; Center for Computational Biology, Johns Hopkins University, Baltimore, MD, USA; Malone Center for Engineering in Healthcare, Johns Hopkins University, MD, USA

**Keywords:** spatially-resolved transcriptomics, quality control, Python, scverse

## Abstract

Spatially-resolved transcriptomics (SRT) generates large and heterogeneous datasets where global (tissue-wide) quality control (QC) metrics often over-aggressively remove biologically meaningful regions or miss localized artifacts. Recently, spatially-aware QC metrics have been introduced in SpotSweeper, but this is limited to the R programming language, which makes it challenging to use these metrics within the Python/scverse ecosystem. Here, we present SpotSweeper-py, a Python equivalent package of SpotSweeper that computes neighborhood-aware *z*-scores for standard QC metrics such as total counts, log total counts, number of detected genes, and percentage of mitochondrial counts. We demonstrate the performance and usability of SpotSweeper-py on two public datasets from the 10x Genomics Visium and VisiumHD platforms. This implementation of local spatially-aware QC metrics enables direct integration with Python/scverse ecosystem, reduces false positives from global quality control while preserving tissue-specific architecture. Plotting utilities are also included for quick visualizations of flagged outliers. By making robust local QC accessible in Python, SpotSweeper-py strengthens the reliability of pipelines for analyzing SRT data. The open-source software is available on PyPI (https://pypi.org/project/spotsweeper).

## 1 Introduction

Spatially-resolved transcriptomics (SRT) technologies have advanced rapidly in recent years, enabling high-throughput mapping of gene expression in complex tissues. Modern SRT platforms include diverse methodologies - from sequencing-based barcoding (e.g. Slide-seq2, 10x Genomics Visium and Visium HD) to imaging-based technologies (e.g. MERFISH, 10x Xenium). Increasingly large and complex spatial datasets are being generated from such technologies [1–3]. These technological innovations provide unprecedented inside into tissue structure and cellular interactions, but they also warrant the need for robust spatially-aware quality control (QC) to ensure reliability and intepretability of downstream analyses.

Historically, computational pipelines for SRT have adopted QC metrics directly from single-cell RNA-seq workflows. Such *global* QC metrics, including total gene expression counts, number of detected genes, and percentage of reads from mitochondrial genes, can be applied across all spatial coordinates (spots or cells) in the tissue to flag low-quality artifacts [4–6]. However, recent work has shown that these global QC metrics may remove biologically meaningful regions as they were not designed for SRT data. In contrast, *local* QC approaches have been developed to address those gaps [7–11]. Notably, SpotSweeper introduced a QC method that utilizes each spot’s neighbors to assess quality in a spatially-aware context [7]. By comparing a given spot’s QC metrics (total counts, gene count, mitochondrial fraction) to the local mean of those metrics in neighboring spots, SpotSweeper flags local outliers (low-quality spots with respect to neighbors) and detects broader regional artifacts that traditional QC pipelines would not detect. This method, validated across SRT technologies, demonstrated its utility and performance over global metrics. However, SpotSweeper is currently implemented as R/Bioconductor package. Meanwhile, the Python ecosystem for single-cell and spatial transcriptomics has been growing rapidly in the past years, driven by libraries such as AnnData, Squidpy, and SpatialData under the scverse ecosystem [12–16]. These powerful tools allow seamless integration of spatial data with major Python workflows for downstream analysis, visualization, and machine learning.

In this manuscript, we bridge the gap by introducing SpotSweeper-py, a Python implementation of the SpotSweeper R/Bioconductor package. SpotSweeper-py is an open-source package available on the Python package index (PyPI) (https://pypi.org/project/spotsweeper), enabling wide adoption and usability to the Python ecosystem. In the following sections, we demonstrate the design of SpotSweeper-py, its integration with major Python pipelines, and a couple use cases across different SRT platforms. By bringing spatially-aware QC into the Python ecosystem, SpotSweeper-py aims to ensure that high-quality data strengthens the robustness of SRT data analysis.

## 2 Use cases of SpotSweeper-py with public SRT datasets

Here, we provide a brief summary of the methodology implemented in SpotSweeper [7] and now SpotSweeper-py. For each spatial coordinate, the method computes a neighborhood-aware robust *z*-score for each spot. Specifically, the method compares the QC metric at a given spatial coordinate to the median of its *k*-nearest neighbors, and scales by the median absolute deviation (MAD) based on all spatial coordinates in the neighborhood. Local outliers are then flagged one or two sided based on user-defined cutoffs. The package directly integrates with AnnData [13] objects in the Python/scverse ecosystem, produces *z*-scores and outlier indicator flags for each spot, and includes plotting utilities both in-line and PDF reporting.

### 2.1 SpotSweeper-Py improves quality control using 10x Genomics Visium data

First, we considered a human breast cancer sample profiled using the 10x Genomics Visium CytAssist FFPE platform [17]. This tissue section had 4,169 spots after loading counts into an `AnnData` [13] object, annotating mitochondrial genes with `MT-`prefixes, and computing standard QC metrics using *Scanpy* [16]. We first considered the log-transformed total UMI counts as the QC metric, where the metric illustrated broad gradients with some tissue gaps and boundary effects typical for a FFPE tissue block (**Figure 1A**). When we consider outliers using a global threshold, specifically the median absolute deviations (MADs) across all spots with 3 MADs, we found that this picks up large and contiguous regions that have systematically low counts, even when those regions are locally homogeneous and plausibly biologically meaningful (**Figure 1B**). In contrast, when we identified outliers using SpotSweeper-py with 36 nearest neighbors and a robust MAD *z*-score cutoff of 3, we found that it revealed small degraded areas (e.g. micro-tears) while mostly retaining surrounding tissue (**Figure 1C**). Specifically, the global (vs local) QC approach (using log total UMI counts) flagged as low-quality 5.49% (229/4169) compared to 1.13% (47/4169) of total spots, respectively. Notably, global QC approaches using the raw (or observed) total counts flagged 0 low-quality outliers in the dataset. This can happen when the raw counts are skewed enough that the global 3 MAD lower-tail threshold falls below all observed values, which is why we recommend analyzing log normalized total counts. The contrast in treatment of regions with systematically low counts captures the core benefits of spatially aware QC.

**Figure 1:**
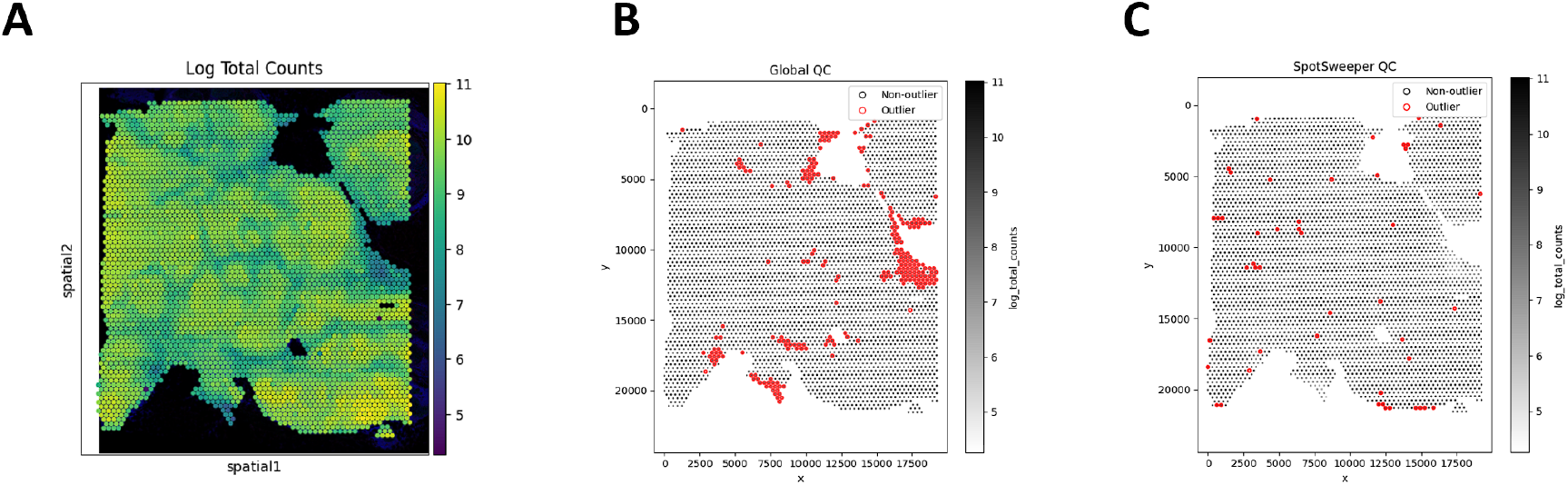
SpotSweeper-py improves quality control using the log-transformed UMI counts as the QC metric. Using a human breast cancer sample profiled using the 10x Genomics Visium CytAssist FFPE platform, **(A)** a spot plot of the log total UMI counts. The black represents bins with zero total UMI counts. **(B)** Detecting low-quality spots global QC thresholds (spots with counts < 3 MADs below the global mean shown in red). **(C)** Detecting low-quality spots using SpotSweeper-py with local QC metrics.

This extends to other QC metrics for SRT data. For example, we see similar patterns of detected outliers across the tissue with the number of detected genes (**Figure 2A**). We found that global QC metrics flagged 3.60% spots (150/4169) as low-quality (**Figure 2B**), compared to 1.92% spots (80/4169) using local QC metrics (**Figure 2C**). Stated another way, global thresholding often discards large regions that are likely biologically meaningful. When using the percent of reads mapping to the mitochondrial genome (**Figure 2D**), we find that global outliers flag as low-quality 14.58% for percent mitochondrial counts (608/4169), which is primarily targeted on the the left region of the tissue, compared to 0.38% (16/4169) of spots flagged as low-quality using local outliers (**Figure 2E-F**). These tend to form small, coherent clusters where mitochondrial content spikes relative to neighbors, corresponding with local technical artifacts.

**Figure 2:**
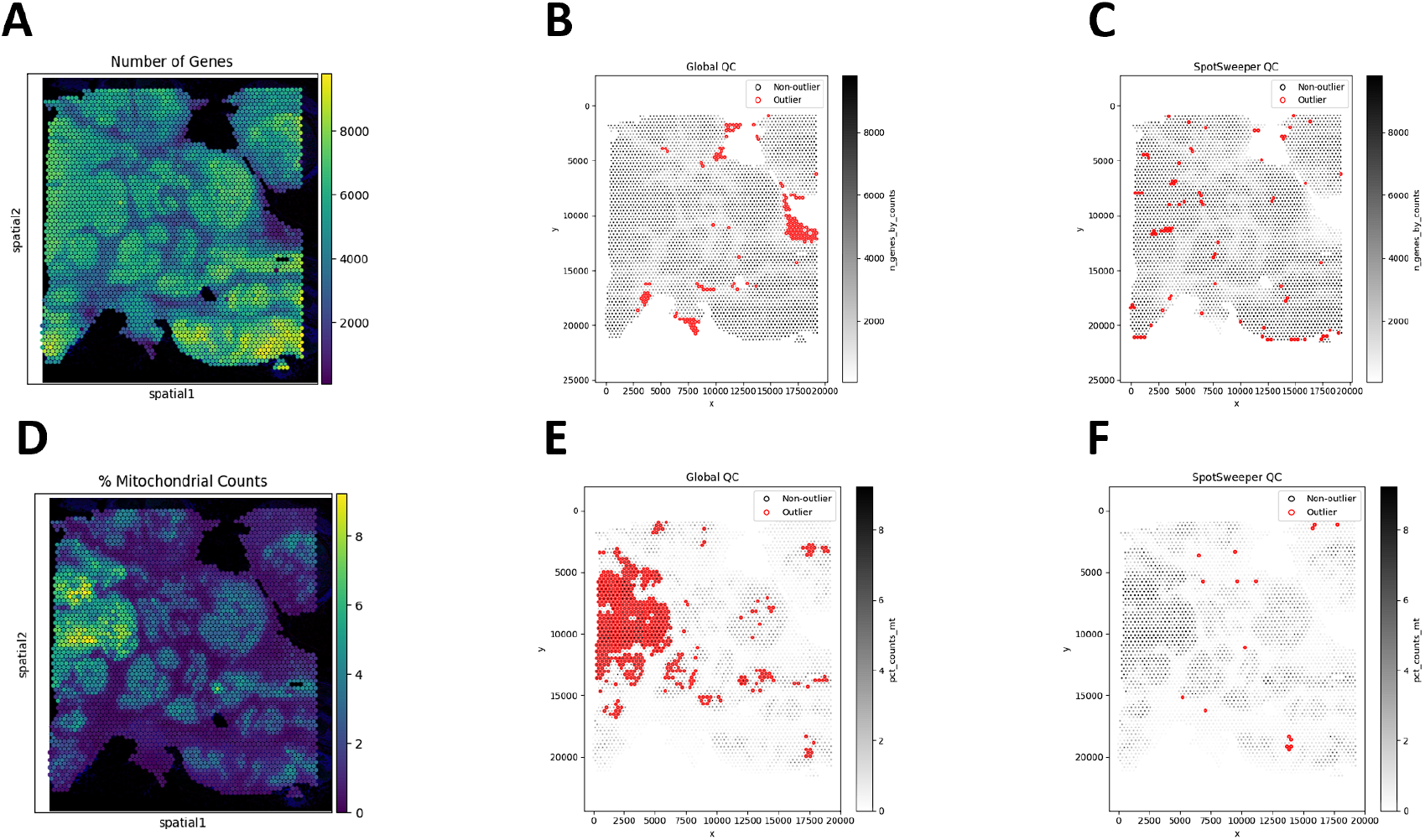
SpotSweeper-py improves quality control using the number of detected genes and the percent of reads mapping to the mitochondrial genome as the QC metric. Similar Figure 1, we used the same human breast cancer sample profiled using the 10x Genomics Visium CytAssist FFPE platform. **(A)** a spot plot of the number of detected genes. **(B)** Detecting low-quality spots global QC thresholds (spots with counts < 3 MADs below the global median shown in red). **(C)** Detecting low-quality spots using SpotSweeper-py with local QC metrics. **(D-F)** Similar to A-C, but using the percent of reads mapping to the mitochrondial genome.

When combining the three QC metrics, we found that there was a marked drop from 20.03% (global) to 2.42% (local) outliers. Similar to our previous findings, we show here that SpotSweeper-py retains biologically meaningful spots[7].

In addition, we examined the distributions of local *z*-scores for standard QC metrics, stratified by whether each spot was flagged as an outlier by either global (**Figure 3A**) or local (**Figure 3B**) QC thresholds (**Table 1**). Using global thresholds, spots labeled as outliers do not consistently correspond to extremes of the local z-score distributions. This reflects the fact that global cutoffs identify spots with extreme absolute QC values, but may fail to detect outlying spots whose deviations are defined relative to their local tissue neighborhood. Conversely, when applying local thresholds, spots flagged as outliers display clear shifts toward the tails of the local *z*-score distributions, demonstrating that the local method captures neighborhood-specific anomalies that are not apparent at the tissue level. Together, these comparisons highlight that local QC metrics identify biologically meaningful, spatially contextualized outliers that global thresholds may overlook and misclassify.

**Table 1:** Outlier rates on the Visium CytAssist FFPE breast cancer sample (*n*=4169). Global: median ±3 MAD (lower-tail for counts/genes; higher-tail for mitochondrial). Local: SpotSweeper-py (*k*=36, cutoff = 3).

| Metric (direction) | Global | Local |
| --- | --- | --- |
| Total counts (lower) | 0.00% (0/4169) | 0.98% (41/4169) |
| Log total counts (lower) | 5.49% (229/4169) | 1.13% (47/4169) |
| Genes detected (lower) | 3.60% (150/4169) | 1.92% (80/4169) |
| % mitochondrial (higher) | 14.58% (608/4169) | 0.38% (16/4169) |
| **Any metric flagged** | **20.03% (835/4169)** | **2.42% (101/4169)** |

**Figure 3:**
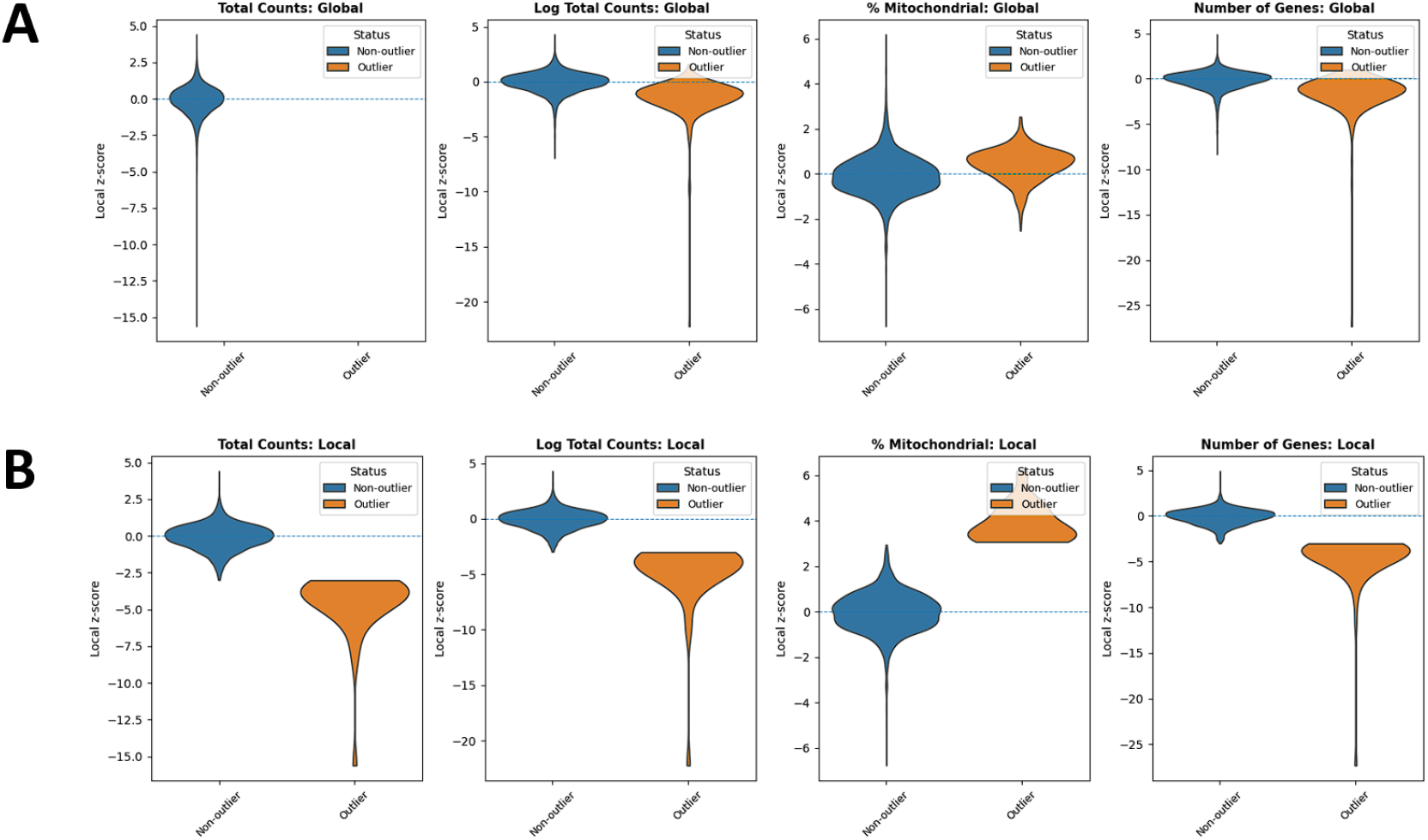
Violin plots comparing outliers and non-outliers flagged by global QC and SpotSweeper QC across multiple QC metrics for a Visium sample. (A) Global QC: total counts, log total counts, percent mitochondrial counts, and number of genes detected. (B) Local QC (SpotSweeper): same metrics with neighborhood-based thresholds. Local *z*-score distributions are shown between outliers (orange) and non-outliers (blue) for each metric.

Importantly, these differences also manifest spatially: spots mis-flagged by global QC form large contiguous regions that SpotSweeper-py successfully preserves, whereas spots uniquely flagged by local QC appear more spatially scattered (**Figure 4**).

**Figure 4:**
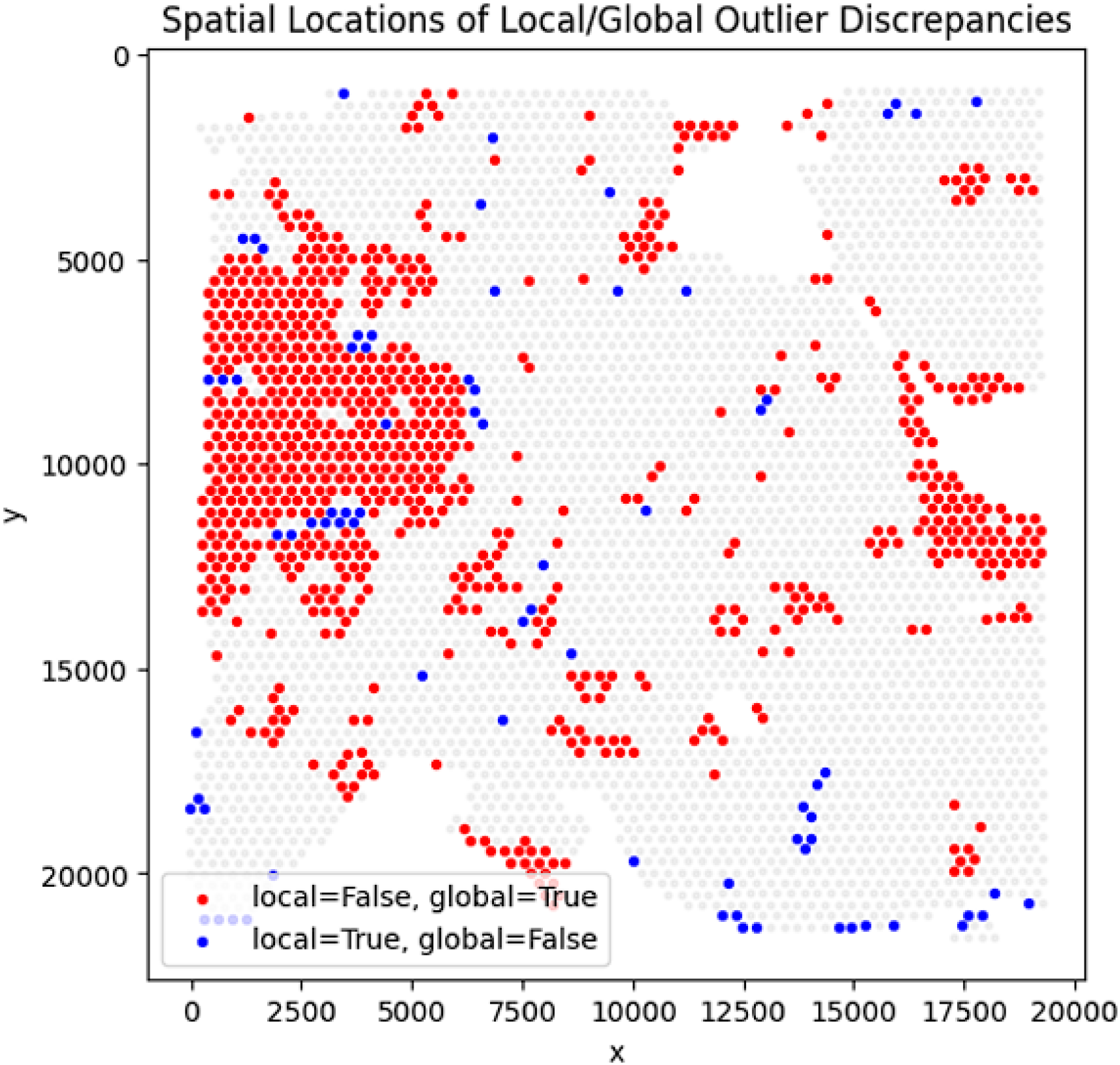
Spatial distribution of discrepancy spots between global and local QC methods for a Visium sample. Red dots mark spots flagged by global QC but not by any local SpotSweeper-py metric, while blue dots mark spots flagged only by local QC. Red discrepancies often appear in contiguous spatial regions, indicating areas that global QC mis-flags but SpotSweeper-py preserves.

### 2.2 SpotSweeper-Py improves quality control using 10x Genomics Visium HD data

We then analyzed a public human breast cancer section profiled with the 10x Genomics VisiumHD CytAssist FFPE platform [18]. Counts were loaded with `spatialdata-io` [15], mitochondrial genes (prefix `MT-`) were annotated, and standard QC metrics were computed with *visiumhd-utils* (built on *Scanpy*) [16]. We primarily used the 8 *µ*m bin level (2 *µ*m results in the next use case). The processed 8 *µ*m `AnnData` object contained *n*=663857 bins. As in the Visium analysis, we contrasted global thresholding (median ±3 MAD; lower-tail for counts/genes; upper-tail for mitochondrial fraction) with SpotSweeper-py using *k*=48 nearest neighbors (from suggestion in SpotSweeper [7]) and a robust *z*-score cutoff of 3. In terms of computational speed, on a CPU node, local outlier detection and plotting for 8 *µ*m total counts completed in 35.7 s, illustrating practical runtimes even at HD scale.

Next, we considered the log-transformed total UMI counts as the QC metric, where the metric exhibits round-shaped structures with localized low-signal areas across the tissue (**Figure 5A**). Using a global threshold, specifically 3 MADs below the global median, we found that outliers cluster in extended low-signal regions and reflect dataset-wide tails rather than neighborhood deviations (**Figure 5B**). In contrast, SpotSweeper-py (with *k*=48 nearest neighbors and a robust MAD *z*-score cutoff of 3) pinpoints small degraded areas while largely retaining surrounding tissue (**Figure 5C**). Specifically, the global approach (using log total UMI counts) flagged as low-quality 2.14% (14211/663857) compared to 1.59% (10551/663857) using the local method. Notably, global thresholds using the raw (observed) total counts flagged 0 low-quality outliers in this dataset (0.00%), whereas the local method flagged 1.06% (7017/663857). This can occur when the raw counts are sufficiently skewed so that the lower-tail threshold falls below all observed values-again the reason we primarily analyze log-total counts.

**Figure 5:**
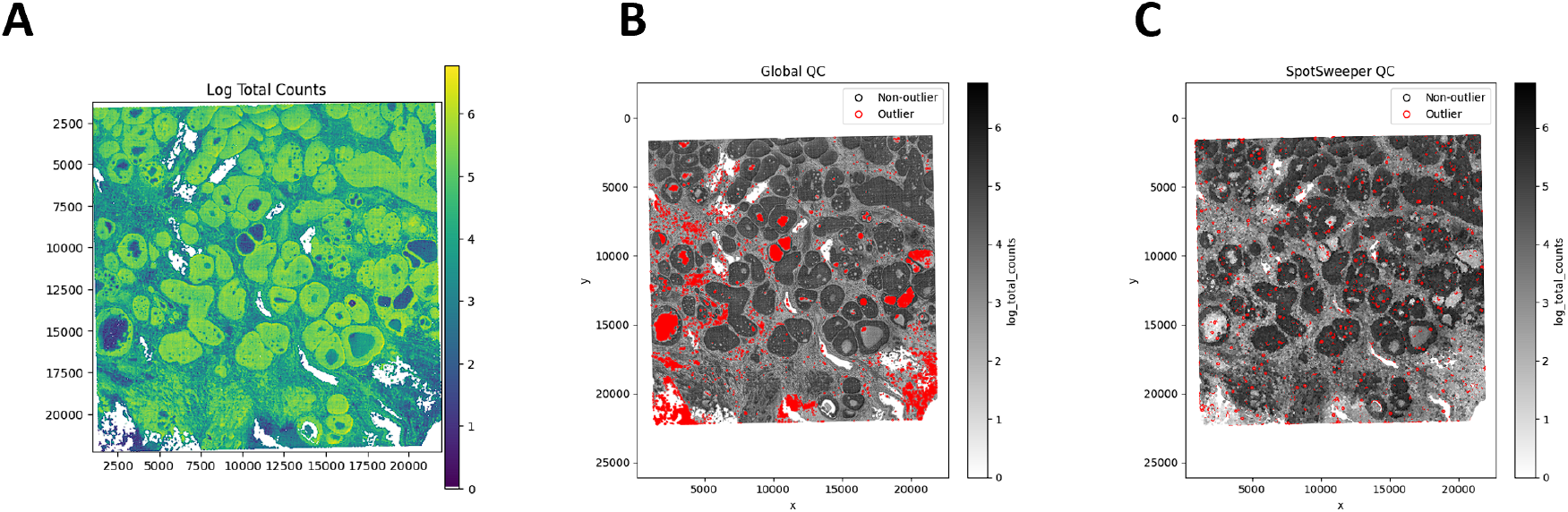
SpotSweeper-py improves quality control using the log-transformed UMI counts as the QC metric on VisiumHD (8 *µ*m). Using a human breast cancer sample profiled using the 10x Genomics Visium CytAssist FFPE platform, **(A)** a bin map of the log total UMI counts. The white represents bins with zero total UMI counts. **(B)** Detecting low-quality bins with a global threshold (bins with metric *<* −3 MADs below the global median shown in red). **(C)** Detecting low-quality bins using SpotSweeper-py with local QC metrics.

The same pattern extends across other QC metrics for SRT data (**Figure 6**). Using the number of detected genes as the QC metric, global thresholds flagged 0.00% (0/663857) of bins (**Figure 6B**), whereas SpotSweeper-py flagged 1.09% (6995/663857), revealing focal anomalies with respect to their neighborhoods (**Figure 6C**). Using the percent of reads mapping to the mitochondrial genome (**Figure 6D**), global thresholds flagged 4.10% (27234/663857) of bins concentrated in broad low-signal regions, compared to 0.22% (1481/663857) flagged as local outliers by SpotSweeper-py (**Figure 6E-F**). These local outliers tend to form small, coherent clusters where mitochondrial content spikes relative to nearby bins, consistent with localized technical artifacts. When combining the three QC metrics, there is a drop from 6.14% (global) to 1.83% (local) of bins flagged as outliers (**Table 2**), reflecting that many globally flagged bins lie in spatially homogeneous low-signal regions that are possibly biologically meaningful regions that spatially aware QC aims to retain.

**Table 2:** Outlier rates on the VisiumHD CytAssist FFPE breast cancer sample at 8um bins (*n*=663857). Global: median ±3 MAD (lower-tail for counts/genes; higher-tail for mitochondrial). Local: SpotSweeperpy (*k*= 48, cutoff = 3).

| Metric (direction) | Global | Local |
| --- | --- | --- |
| Total counts (lower) | 0.00% (0/663857) | 1.06% (7017/663857) |
| Log total counts (lower) | 2.14% (14211/663857) | 1.59% (10551/663857) |
| Genes detected (lower) | 0.00% (0/663857) | 1.09% (6995/663857) |
| % mitochondrial (higher) | 4.10% (27234/663857) | 0.22% (1481/663857) |
| **Any metric flagged** | **6.14% (40754/663857)** | **1.83% (12173/663857)** |

**Figure 6:**
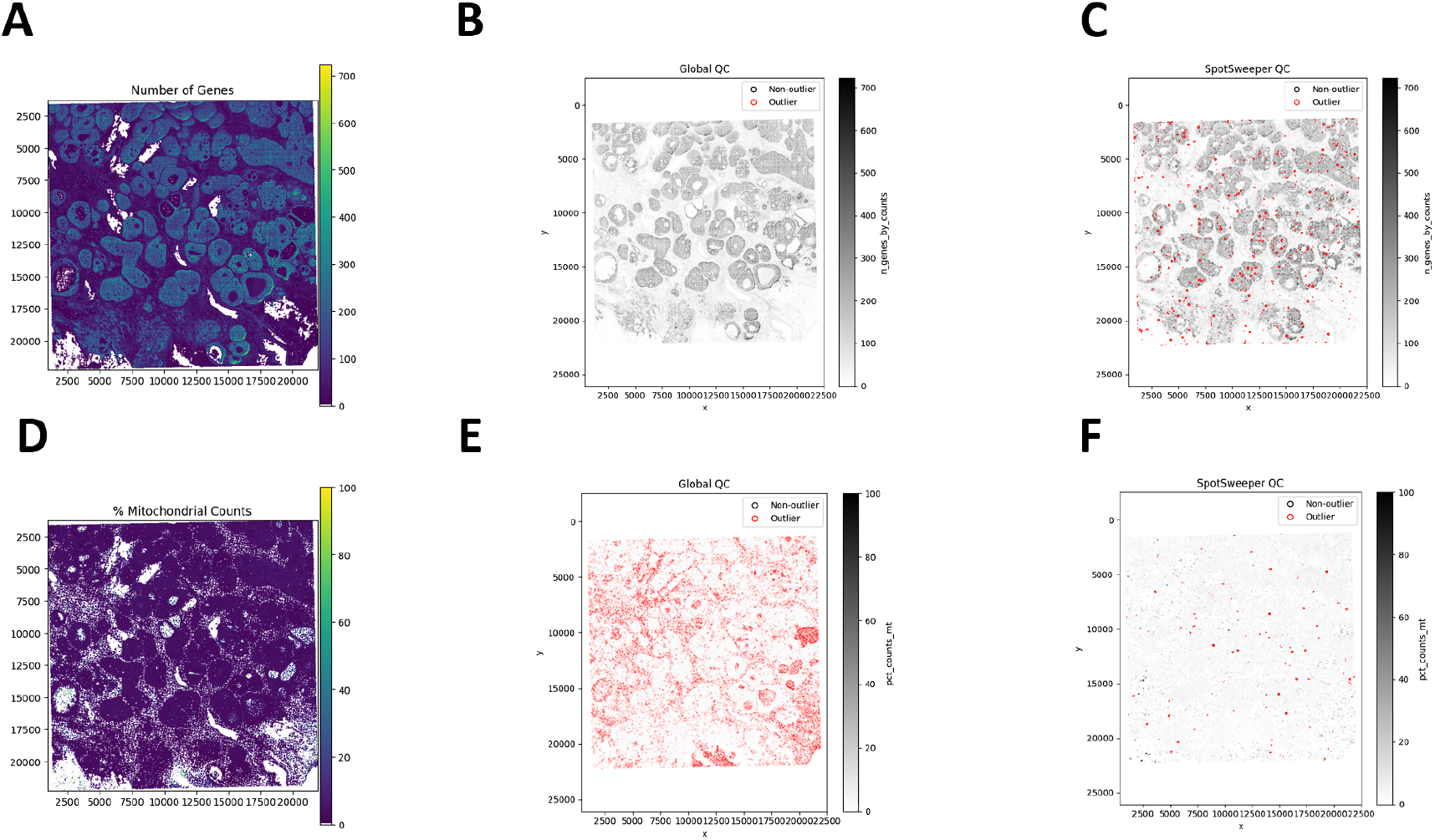
SpotSweeper-py improve quality control using the number of detected genes and the percent of reads mapping to the mitochondrial genome as the QC metric. Similar to Figure 5, we used the same human breast cancer sample profiled using the 10x Genomics Visium HD CytAssist FFPE platform (8 *µ*m resolution). **(A)** a spot plot of the number of detected genes. **(B)** Detecting low-quality spots global QC thresholds (spots with counts < 3 MADs below the global mean shown in red). **(C)** Detecting low-quality spots using SpotSweeper-py with local QC metrics. **(D-F)** Similar to A-C, but using the percent of reads mapping to the mitochrondial genome.

In addition, we compared the distributions of local *z*-scores for QC metrics stratified by whether a bin was flagged as an outlier under either global (**Figure 7A**) or local (**Figure 7B**) thresholds (**Table 2**). When applying global thresholds, non-outlier and outlier groups show limited separation in their local *z*-score distributions for several metrics, and for total counts and number of detected genes, no global outliers were detected at all. This indicates that global criteria primarily capture extreme absolute QC values and may overlook spatially localized anomalies. In contrast, under local thresholds, bins flagged as outliers show clear shifts toward the extremes of the local *z*-score distributions across all metrics, demonstrating that the local method effectively identifies neighborhood-relative deviations that are not apparent when using global cutoffs.

**Figure 7:**
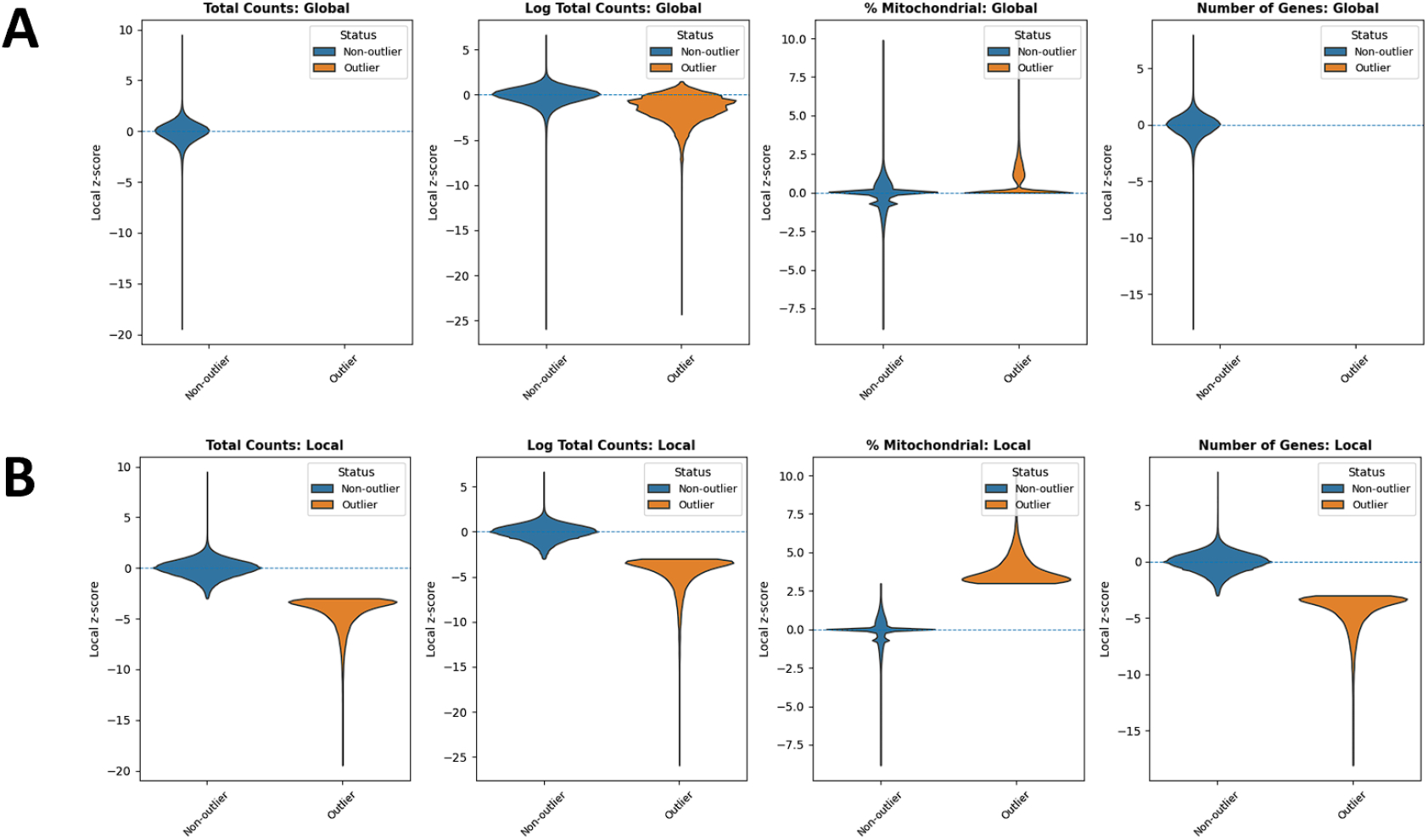
Violin plots comparing outliers and non-outliers flagged by global QC and SpotSweeper QC across multiple QC metrics for a Visium HD sample profiled at 8 *µ*m bin resolution. (A) Global QC: total counts, log total counts, percent mitochondrial counts, and number of genes detected. (B) Local QC (SpotSweeper): same metrics with neighborhood-based thresholds. Local *z*-score distributions are shown between outliers (orange) and non-outliers (blue) for each metric.

### 2.3 Extended Use Case: SpotSweeper-Py improves quality control using 10x Genomics Visium HD data profiled at 2*µ*m level

Finally, we evaluated SpotSweeper-py on the highest–resolution setting of the Visium HD CytAssist FFPE platform, using the 2*µ*m bin representation of the same data analyzed above. After loading counts using `spatialdata-io` and computing standard QC metrics with *visiumhd-utils*, the resulting `AnnData` object contained *n*=10535119 bins. As with the 8*µ*m analysis, we compared global median ±3 MAD thresholds to SpotSweeper-py with *k*=48 nearest neighbors and a robust local *z*-score cutoff of 3. Despite the extremely large bin count, SpotSweeper-py completed local QC computations and visualization in under 15 minutes on a standard CPU node, demonstrating that neighborhood-based QC remains computationally feasible even at full Visium HD resolution.

Figure 8. demonstrates that SpotSweeper-py remains effective even at the highest resolution of the Visium HD platform. Despite the greatly increased bin count, local QC continues to isolate small, spatially coherent defects without over-flagging large biologically meaningful regions. Across QC metrics (log total counts, number of genes, percent mitochondrial counts), SpotSweeper-py consistently identifies compact neighborhoods of outlier bins while preserving the majority of intact tissue.

**Figure 8:**
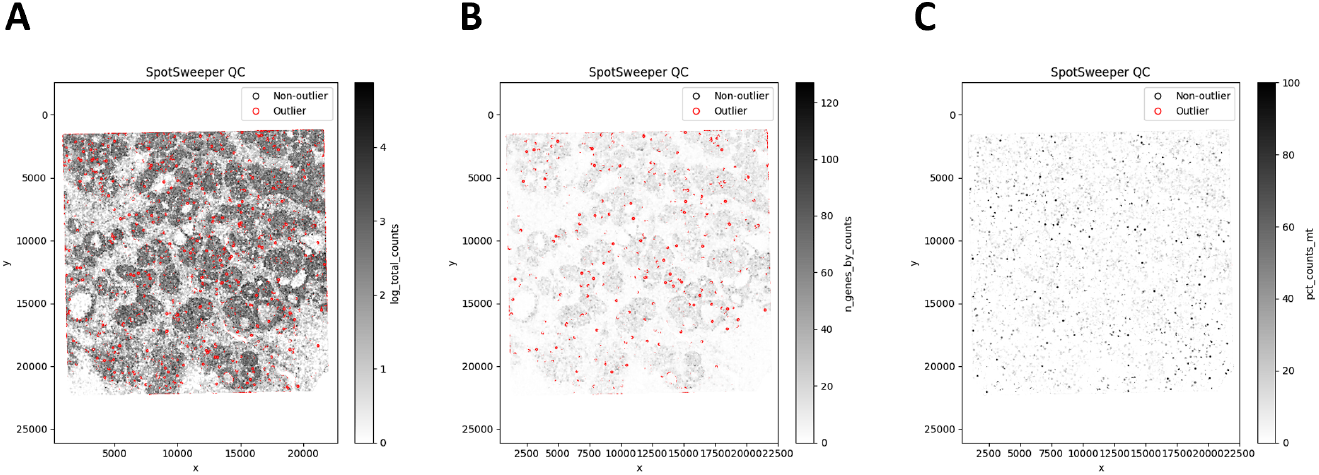
Outputs of SpotSweeper QC on a Visium HD tissue section profiled at 2 um resolution. Outlier spots (red) are identified using local thresholds (48 neighbors). (A) Low outliers based on log total UMI counts. (B) Low outliers based on number of genes detected. (C) High outliers based on percent mitochondrial counts.

In contrast to the outlier maps, the distributional behavior of QC metrics at 2*µ*m resolution further highlights the limitations of global thresholding. Under global QC, only the percent mitochondrial metric produced any outliers at all: total counts, log total counts, and number of detected genes showed no bins exceeding global 3-MAD cutoffs (**Figure 9A**). When examining the local *z*-score distributions (**Figure 9B**), bins identified by SpotSweeper-py show clear and consistent shifts toward the tails of their respective local *z*-score distributions. This separation demonstrates that local QC captures spatially aware deviations that define technical artifacts at a high resolution, whereas global QC merely flags significant global shifts.

**Figure 9:**
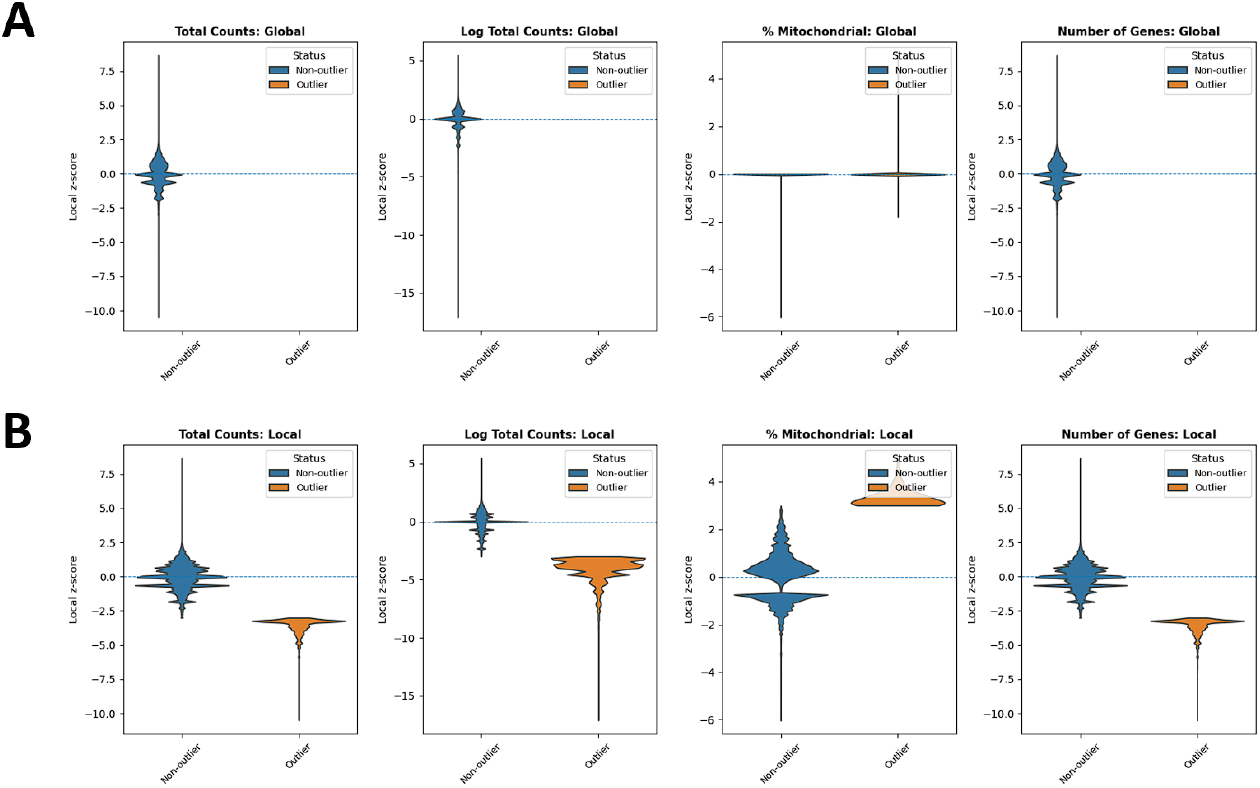
Violin plots comparing outliers and non-outliers flagged by global QC and SpotSweeper QC across multiple QC metrics for a Visium HD sample profiled at 2um bin resolution. (A) Global QC: total counts, log total counts, percent mitochondrial counts, and number of genes detected. (B) Local QC (SpotSweeper): same metrics with neighborhood-based thresholds. Local *z*-score distributions are shown between outliers (orange) and non-outliers (blue) for each metric.

## 3 Discussion

Spatial transcriptomics data exhibit heterogeneity driven by both technical and biological factors, such as tissue boundaries, gradients, and tissue degradation. Therefore, applying global, dataset-wide thresholding to quality control metrics could remove the entirety of biologically significant regions or miss regions with small but important tissue defects. Our PyPI package, SpotSweeper-py, addresses this issue by building a spatially-aware quality control framework that identifies outliers based on local robust *z*-scores that integrates seamlessly into the Python/scverse ecosystem. When conducting analyses on datasets from two sample spot-based spatial transcriptomics technologies, Visium and Visium HD (at 8*µ*m level), SpotSweeper-py reduced false positives with respect to outliers identified using global MAD thresholds and revealed small, spatially-aware artifacts consistent with biological structure. As the implementation of SpotSweeper-py integrates with scverse, SpatialData, and AnnData ecosystem in Python, the annotations for local outliers could easily transfer to downstream steps like clustering and differential expression. As a result, the reliability and interpretability of spatial transcriptomics data analyses could be improved in Python-based workflows.

However, there are still some limitations to consider. SpotSweeper-py assumes accurate spatial coordinates and relatively uniform capture spot density. Any technical issues such as mis-stitching could significantly bias local statistics and may impact the performance of our algorithm by outputting false positives. It is also important to note that run-time and memory usage scales up as the number of spots and bins increases due to *k*-nearest neighbor construction. In our example use cases, the algorithm is feasible on a CPU - the runtime for a Visium section and a Visium HD section profiled on 8*µ*m bins are under 1 minute. However, the runtime quickly scales up for the same Visium HD section, profiled at 2*µ*m bins. Tshe run-time could take up to 15 minutes for outlier calculations in that setting. Finally, the local outliers detected from SpotSweeper are indicators that solely based on sequencing metrics: additional histology evidence is also recommended before filtering.

Here, we demonstrated how to control bias in outlier detection using SpotSweeper-py. Robust and neighborhood-aware statistics are computed based on the median and MAD, thus limiting the influence of extreme values and heavy-tailed distributions. Log(1+x) transforms are performed on UMI counts before conducting local outlier analysis in order to stabilize variance for count metrics. We also recommend investigating common outliers from all 4 QC metrics to reduce dependence on one particular metric.

Looking to the future, SpotSweeper-py will be extended from just local outlier detection to include broader spatial artifact detection as implemented in SpotSweeper [7], mirroring the functionalities in the implementation in R. As a result, we could expect additional identification of contiguous and systematic low-quality regions. Additional extension directions include multi-scale QC that includes outliers across neighborhood radii, dynamic neighbor number adjustment based on the shape of data, as well as additional interactive visualization and data reporting. Collectively, these developments aim to make local and spatially-aware QC a transparent and standard component of spatial transcriptomics analysis pipelines and therefore strengthen the credibility of downstream results.

## 4 Methods

### 4.1 Implementation of SpotSweeper-py

#### 4.1.1 Overview and relationship to SpotSweeper (R)

Our package, `spotsweeper_py` re-implements, in Python, the local QC outliers detection and plotting functionalities of the original SpotSweeper R package, which introduced spatially aware QC to detect both local outliers and regional artifacts in spatial transcriptomics outputs using standard QC metrics (e.g. library size, detected genes, mitochondrial fraction). Here, we focus on the local outlier component (parallel to `localOutliers()` function in the R package) and on plotting utilities. Our proposed Python package emphasizes a clean and lightweight Python API for local outlier detection and publication-ready visualization that integrates directly with major downstream workflows like `AnnData/ Scanpy`, providing a Python-native option.

#### 4.1.2 Essential data inputs

The core function to detect local outliers,

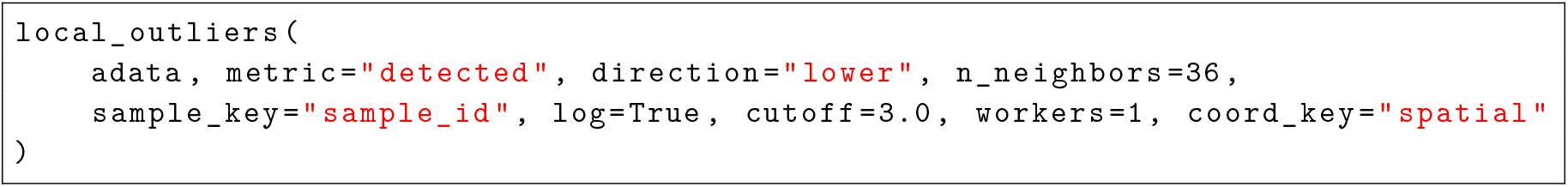

operates on a single `anndata.AnnData` object and expects the following components to be available in the `AnnData` object.

- Spatial Coordinates: `adata.obsm[coord_key]` (default: “spatial”) with shape (n_spots * 2).
- Sample Identifier: `adata.obsm[sample_key]` (default: “sample_id”) to ensure neighbors are computed within independent samples.
- QC Metrics: `adata.obs[metric]` (default: “detected”). This could be substituted with other columns, for instance total counts, log total counts, or mitochondrial ratio.

Strict validity checks are performed on these columns. If the columns are missing, a `KeyError` will be raised. For invalid parameters in addition to those columns, a `ValueError` will be raised, preventing the program from running. Other parameters will be explained in sections below.

#### 4.1.3 Neighborhood construction

For each sample, the local outliers function learns a *k*-nearest neighbor (kNN) structure over spatial coordinates with `scikit-learn`’s nearest neighbor function [19]. We use the “auto” algorithm in *scikit-learn* package to let it choose the fastest model for different data samples. The algorithm will select between a tree-based or brute force strategy depending on data size and dimensionality. Neighbor indices are then returned using scikit-learn’s helper functions. Typical defaults use 36 as the neighborhood size (equivalent with the R version, generalizable across many spatial transcriptomics technologies), but this is modifiable via n_neighbors parameter in the local outliers function. There is also an opportunity for parallelization, to be controlled via workers parameter. It defaults to 1, which means no parallelization in computing.

#### 4.1.4 Robust local *z*-score calculations and outlier determination

Let *x*_*i*_ be the (optionally log-transformed) QC value at spot *i*, and let *N* (*i*) denote the indices of its *k* spatial neighbors. We compute a modified/robust *z*-score:

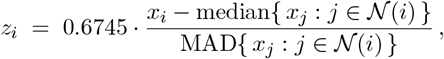

where MAD, the median absolute deviation, takes the following form: MAD(*S*) = median_*j*∈*N* (*i*)_ | *x*_*j*_ − median{*x*_ℓ_ : ℓ ∈ *N* (*i*)}. If MAD is zero or non-finite (when we have degenerate neighborhoods), we conservatively set the robust *z*-score to be 0. Non-finite results are also handled and set to 0. Multiple parameters in the local outlier functions control the flagging process, including direction, which determines whether flagging is one-sided (“higher” or “lower”), or two-sided (“both”) along with:

- `higher`: will flag outliers if *z*_*i*_ *>* cutoff,
- `lower`: will flag outliers if *z*_*i*_ *<* −cutoff,
- `both`: two-sided, will flag outliers if |*z*_*i*_| *>* cutoff.

The default `cutoff = 3.0` is conservative and corresponds to classic robust *z*-score practice. Both the cutoff value and direction could be modified via function parameters.

#### 4.1.5 Log transform and processing with AnnData objects

If the log parameter is set to True (by default), we apply log(1 + *x*) to reduce the impact of extremely large outliers. We write the log-transformed results into a new column `{metric}_log in adata.obs`. Note we do not overwrite the column if it already exists in `adata.obs.` This process ensures reproducibility of the pre-processing pipeline.

#### 4.1.6 Local outlier function output

The function returns the updated `AnnData` object, with the following updated columns in adata.obs:

- `adata.obs[f”{metric}_z”] (float)`: stores per-spot robust *z*-scores.
- `adata.obs[f”{metric}_outliers”] (bool)`: stores whether the spot is flagged by the selected outlier direction and *z*-score cutoff.

These columns are index-aligned with the input AnnData object.

#### 4.1.7 Additional visualization utilities

In addition to the main local outlier functions, we included two extra helper modules for visualizing the location of local outliers in a spatial context. The corresponding functions are:

- `plot_qc_metrics`: returns a scatterplot of single sample spatial coordinates, with continuous color for the metric, and red borders for identified outliers. We set an equal aspect ratio, invert the y-axis to match conventional tissue orientation, and create the colormap via `LinearSegmentedColormap` using either default colors, or user-supplied colors.
- `plot_qc_pdf`: similar to the previous plotting function, while writing all output figures to a multipage PDF with customizable point size and figure dimensions.
- **Function validation and tests via** `pytest`

We have setup a `pytest` suite for the package. It verifies:

- exact robust *z*-score calculations on synthetic inputs.
- creation, correct length and type of newly created columns in `adata`.
- correct detection of an extreme outlier.
- correct log-transformation and column creation.
- correct error-handling for all cases.
- successful generation of a PDF in a temporary directory.

Those tests ensure the local outlier function as well as the plotting functions could produce accurate and defensible outputs.

### 4.2 Operation

#### 4.2.1 Minimal system requirements

- Software: Python, preferably has a version newer than 3.9.
- Operating System: Linux, Windows, or MacOS.
- Hardware: CPU is enough to run the local outlier functions (no GPU required). The usage of GPU could be helpful for multi-sample analyses, or for technologies like VisiumHD, where we observe millions of bins.

#### 4.2.2 Run-time package dependencies

The packages should appear standard to major Python downstream analysis workflows: Core: `anndata, numpy, pandas, scikit-learn, matplotlib`.

Testing: `pytest`

#### 4.2.3 Workflow overview

To start the pipeline, we load and prepare the datasets, in the form of `AnnData` objects. Specifically, we need to ensure that `adata.obsm[“spatial”], adata.obs[“sample_id”]` and a QC metric column (ex. `adata.obs[“detected”]`) are present in the `AnnData` object.

Then, we call the local outlier function to detect outliers.

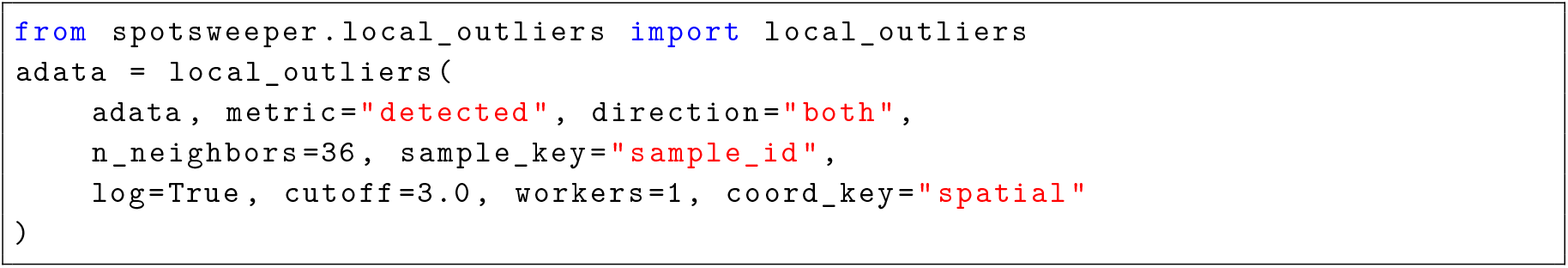

This snippet annotates `detected_z` and `detected_outliers` in `adata.obs`.

We could then use the boolean outlier column `detected_outliers` to flag or filter local outlier spots before further downstream processing such as normalization, clustering, or differential expression analysis.

Optionally, we could visualize the result of local outlier detection:

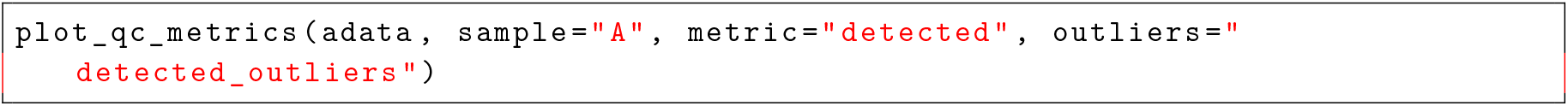

This function could interactively show the figures in a Jupyter notebook, or return a plot of the sample as well as flagged outliers.

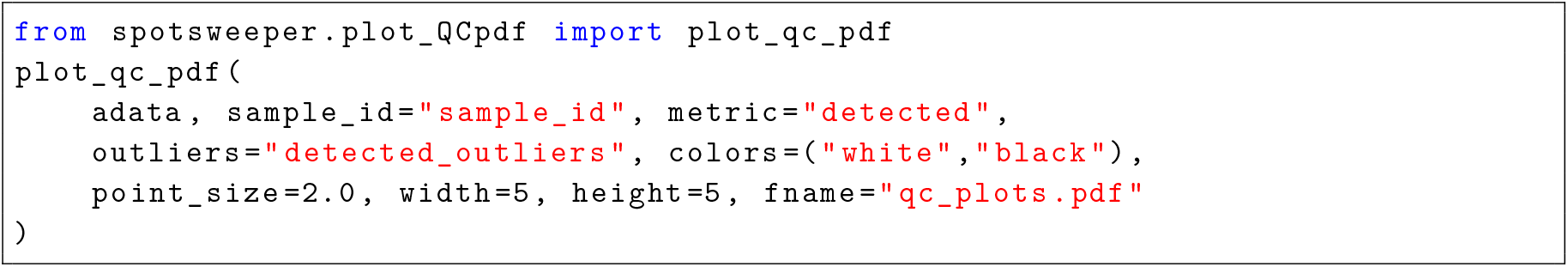

Or, we could take advantage of `plot_qc_pdf` function to generate the plots on a PDF.

#### 4.2.4 Parameter guidance for typical workflows

There are a couple adjustable parameters in the local outlier functions. Generally, we make the following recommendations:

- Neighbors (`n_neighbors`): start with 24–48. Smaller *k* sharpens locality but increases variance; larger *k* stabilizes estimates but may dilute local variance.
- Direction (`direction`): “`lower`” for metrics where extremely low values imply poor quality (for example, number of detected genes), “`higher`” for metrics like mitochondrial fraction, “`both`” if either tail is undesirable.
- Threshold (`cutoff`): default 3.0; consider 2.5 or lower when the tissue is complex, and more outliers need to be considered even with low confidence.
- Log transform (log): recommended `True` for heavy-tailed metrics.

### 4.3 Data availability

The datasets used for the analyses in this manuscript can be downloaded from https://www.10xgenomics.com/datasets/gene-and-protein-expression-library-of-human-breast-cancer-cytassist-ffpe-2-standard (Visium) and https://www.10xgenomics.com/datasets/visium-hd-cytassist-gene-expression-libraries-human-breast-cancer-ffpe-if (Visium HD).

### 4.4 Software availability

SpotSweeper-py is an open-source software on PyPI (https://pypi.org/project/spotsweeper). Code to reproduce all preprocessing, analyses, and figures in this manuscript is available from GitHub at https://github.com/danielchen05/SpotSweeper_py_paper. We used SpotSweeper-py version 0.2.6 for the analyses in this manuscript. Analyses were performed with Python version 3.9.

## Acknowledgments

We thank the maintainers of the Joint High Performance Computing Exchange (JHPCE) compute cluster at Johns Hopkins Bloomberg School of Public Health for providing essential computing resources. We also thank Hicks lab for their constructive comments and suggestions.

## Author Contributions Statement

Visium and VisiumHD data were processed and analyzed by XC. XC and SCH conceptualized the SpotSweeper-py extension of the original SpotSweeper package. The formal analysis and benchmarking was led by XC with suggestions from SCH and MT. Figures were designed by XC. Software was developed by XC with input from other co-authors. SCH administered and supervised the project. XC and SCH wrote the original draft of the text and all co-authors edited and approved the final manuscript.

